# The Establishment of Transgenerational Epigenetic Inheritance in the *C. elegans* Germline is Mediated by Lipid Metabolism

**DOI:** 10.1101/2020.11.04.367854

**Authors:** Dan Peng, Chen Wang, Kai-Le Li, Zhi-Xue Gan, Yong-Hao Li, Hao-Wei Wang, Qiu-Yu Li, Xi-Wei Liu, He-Yuan Sun, Yuan-Ya Jing, Qiang Fang, Qian Zhao, Lei Zhang, Huan-Huan Chen, Hui-Min Wei, Jin Sun, Huan-Yin Tang, Xiao-Mei Yang, Jiang-Feng Chang, Feng Sun, Ci-Zhong Jiang, Hong-Bin Yuan, Wei Li, Fang-Lin Sun

## Abstract

Environmental stress-induced epigenetic changes are inherited by germlines and profoundly affect the behavior and physiology of subsequent generations. However, the mechanisms by which acquired transgenerational epigenetic imprints are established in the parental germline remain poorly understood. In this study, we used *Caenorhabditis elegans* as a model system and demonstrated that a key node of metabolism-Pod-2, an acetyl-CoA carboxylase, together with vitellogenins (Vits), cholesterol-binding/transport proteins, guide establishment of the acquired transgenerational epigenetic modifications H3K27me3 in the parental germline. The loss of function of a Vit; its transporter or receptor; or its binding partner, pod-2, resulted in the loss of acquired epigenetic memory in the germline. Our findings indicate that lipid metabolism is a critical mediator for establishing transgenerational epigenetic imprints in the germline.

## MAIN TEXT

Experiences of adverse environmental stress, e.g., fear-inducing conditions, drugs, diet and endocrine disruption, or chronic exposure to a pathogenic bacterium, are known to affect the behavior and physiology of offspring^1–4^. The inheritance of acquired traits in subsequent generations is believed to profoundly impact the survival of animals under fluctuating environmental conditions^5, 6^ Such transgenerational characteristics acquired from ancestral experiences also have a significant influence on animal health in successive generations of offspring. Altered epigenetic imprints/regulators, such as the modification of histones or DNA/RNA and small RNAs, are crucial in the trans- or intergenerational inheritance of characters through ancestral germlines^3–5, 7–11^. However, the mechanism by which the transgenerationally acquired inheritable epigenetic memory is initially established in the germline of parental animals remains poorly understood.

We used *Caenorhabditis elegans* as a model system to address this problem. Chronic exposure to the pathogenic bacterium *Pseudomonas aeruginosa* PA14 modulates behavioral and cognitive functions in worms for multiple generations^1, 2, 4^. To obtain a more consistent transgenerational behavior assay, we modified the training process by hatching eggs on one side of a barrier lawn of the pathogenic bacterium *P. aeruginosa* PA14 with passive exposure to the pathogen for 72 hours (Fig. 1a). The olfactory assay demonstrated that animals trained with *P. aeruginosa* PA14 for 72 hours were significantly more likely to migrate away from *P. aeruginosa* PA14 than the naïve group for up to four generations: P0, F1, F2 and F3 (Fig. 1b). This condition was thus chosen for further studies.

**Fig. 1.**
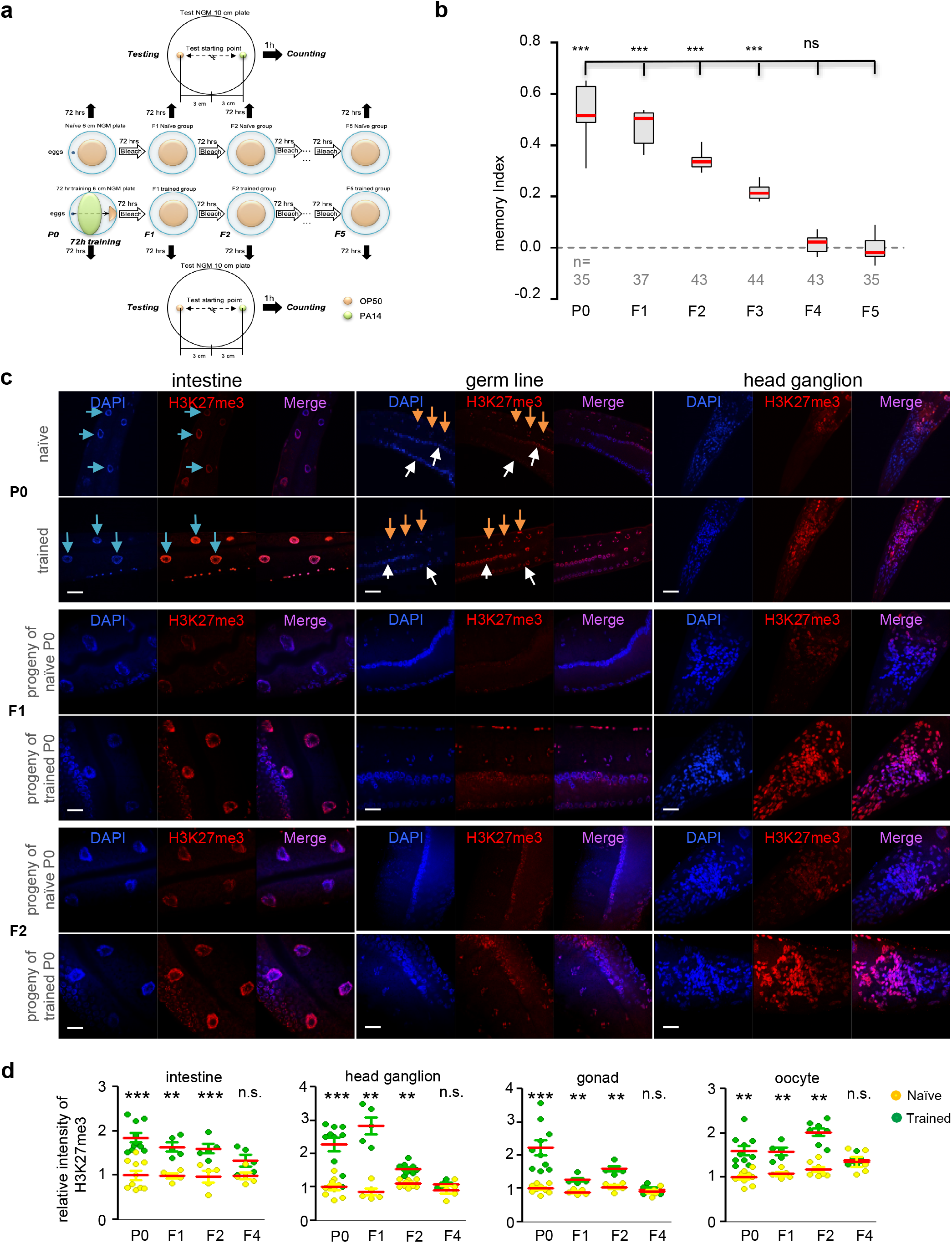
H3K27me3 was identified as a hallmark for pathogenic bacterium-induced parental germline and transgenerational inheritance. **a**, Schematic illustration of the bacterial choice assay, 72-hour training protocol, and multigenerational testing protocol. **b**, Memory index of P0 worms after 72 hours of training on *P. aeruginosa* PA14 and their F1-F5 offspring. n, number of independent assays, approximately 100 worms/assay; p-values were generated by one-way ANOVA with the Dunnett correction (n.s., not significant; *p < 0.05, **p < 0.01, and ***p < 0.001). **c**, Illustration showing that *P. aeruginosa* PA14 increased the levels of global histone H3K27me3 in the intestine, germline and head ganglion in P0 adult animals and their F1 and F2 descendants but not in their F4 descendants. Representative worms were fixed and immunostained with 4’,6-diamidino-2-phenylindole (DAPI) (blue) and antibody against H3K27me3 (anti-H3K27me3) (red). The merged signals are shown in pink. Blue, white and yellow arrows indicate the intestine, gonad and oocyte, respectively. **d**, Quantification of H3K27me3 in the intestine, head ganglion, gonad and oocyte in P0 adult animals. n≥5 worms analyzed, means ± SEMs; p-values were generated by one-way ANOVA with the Dunnett correction (n.s., not significant; *p < 0.05, **p < 0.01, and ***p < 0.001).

The next step was defining a specific acquired transgenerational epigenetic marker/imprint after exposure to the pathogenic bacterium *P. aeruginosa* PA14/stress in worms. Immunofluorescent staining at the whole-animal level was utilized. The trained P0, F1, F2, F3 and F4 worms were separately fixed and immunostained with antibodies against the histone modifications H3K27me3 and H3K9me2/3, markers of repressive transcription, and H3K27ac and H4K16ac, markers of active chromatin^7, 12^. Compared to the naïve worms, all of the *P. aeruginosa* PA14-trained worms exhibited an aberrant increase in H3K27me3 in various tissues, including the germline (gonad and oocytes), intestine, and head ganglion, from parental animals (P0) to F3 animals (Fig. 1c, d). The accumulation of H3K27me3 was accompanied by a robust decrease in H3K27ac (Extended Data Fig. 1a, b), an antagonizing epigenetic marker of H3K27me3^12^, in the corresponding organs of the animals, as anticipated. Thus, these results support that H3K27me3 and H3K27ac serve as transgenerational epigenetic markers for *C. elegans* trained with the pathogenic bacterium *P. aeruginosa* PA14.

To explore whether dynamics in the acquired epigenetic changes H3K27me3 and H3K27ac are linked to avoidance behavior in the *P. aeruginosa* PA14-trained worms, we treated the animals with GSK343, a chemical inhibitor of EZH2^12, 13^, which encodes the histone methyltransferase for H3K27me3, followed by TSA, an inhibitor of histone deacetylases (HDACs), at different concentrations. The presence of both GSK343 and TSA resulted in a significant loss in the behavioral memory of the trained P0 animals in a dose-dependent manner (Fig. 2a, b). Worms expressing mutant MES-2/mes-2, an ortholog of mammalian EZH2, and an HDAC ortholog also showed a similar loss of acquired memory (Fig. 2c). These results further support the notion that an acquired increase in H3K27me3 or decrease in H3K27ac served as an ideal transgenerational epigenetic marker in *C. elegans* under pathogenic conditions.

**Fig. 2.**
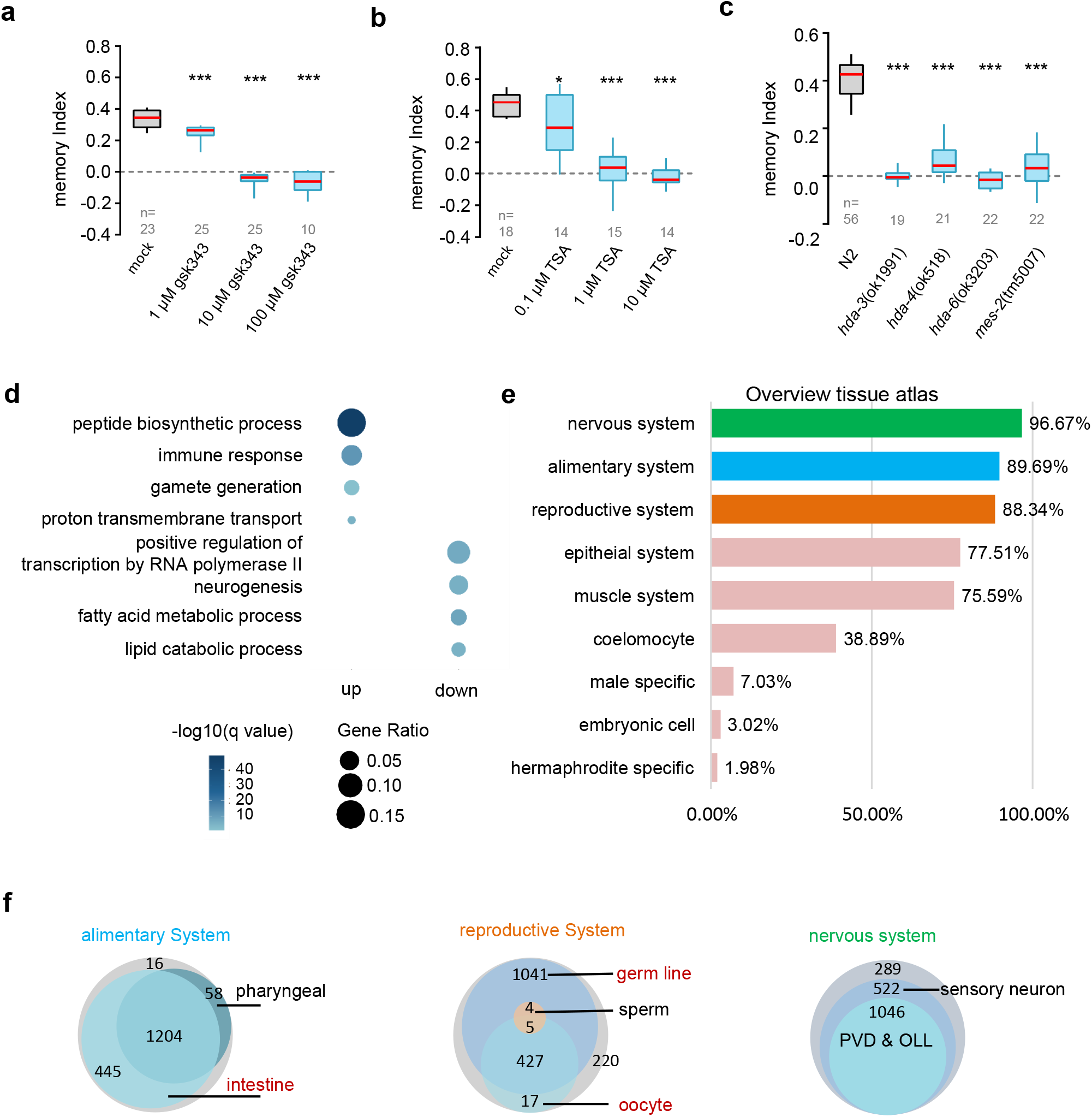
Screening of inhibitors and epigenetic regulators of H3K27me3 in transgenerational memory inheritance. **a**, The presence of the EZH2-specific inhibitor GSK343 blocked *P. aeruginosa* PA14-induced avoidance behavior. **b**, The presence of the histone deacetylase (HDAC) inhibitor trichostatin A (TSA) blocked *P. aeruginosa* PA14-induced avoidance behavior. **c**, PA14 failed to induce avoidance memory in animals with mutations in HDAC family members and the histone H3K27me3 methylase EZH2/mes-2. The number (n) of animals used for the test and the concentrations of the inhibitors are indicated; p-values were generated by one-way ANOVA with the Dunnett correction (n.s., not significant; *p < 0.05, **p < 0.01, and ***p < 0.001). **d**, Dot plot of selected biological process GO terms enriched in genes found by the Wald test to be differentially expressed genes (DEGs) (q-value < 0.1) between N2 *P. aeruginosa* PA14-trained worms compared to N2 naïve worms, as measured by RNA-seq (DEseq). The size of the dot represents the ratio of genes enriched in the specific function. The color gradient represents the FDR q-value forenrichment of the term in the form of a negative logarithm. **e**, Bar plot of the overview tissue atlas showing the percentage of reported tissue expression among the DEGs. The anatomical terms on the y-axis from the database were manually categorized into nine groups. **f**, The Venn diagram on the left indicates the proportion of DEGs expressed in the intestine and pharynx over all reproductive system DEGs. The middle Venn diagram indicates the proportion of DEGs expressed in the germline, oocytes and sperm over all reproductive system DEGs. The Venn diagram on the right indicates the proportion of DEGs expressed in the nervous system.

To understand the mechanism leading to the aberrant change in H3K27me3 and H3K27ac in the germline of the trained animals, an RNA-seq analysis was performed using total RNA extracted from the *P. aeruginosa* PA14-trained and naïve animals at P0. The presence of *P. aeruginosa* PA14 led to the altered transcription of 1215 upregulated and 1154 downregulated target genes (Fig. 2d). However, among the affected genes, we failed to identify any known histone modification enzymes that directly target H3K27me3 or H3K27ac. A large number of upregulated/downregulated genes are associated with the immune response, the peptide biosynthetic process, fatty acid metabolism, and lipid catabolic processes (Fig. 2d). Gene ontology (GO) term analysis further demonstrated that the functions of the top three clusters of affected genes are associated with the neurological, alimentary and reproductive systems (Fig. 2e, f).

Vitellogenins (Vits), a family of yolk proteins encoded by six Vit genes, are known to be highly expressed in the adult worm intestine^14^. In *C. elegans,* the synthesized yolk proteins secreted from the intestine into the body cavity are taken up by the gonad and then reach the oocyte^15–17^. We therefore examined whether Vits/yolk proteins affect establishment of the transgenerational epigenetic modifications H3K27me3 and H3K27ac in the germline of *P. aeruginosa* PA14-trained parental animals. We used mutants of the Vit *vit-1* and the Vit transporter *nrf-6—*to perform immunofluorescence (IF) experiments (Fig. 3a). The level of H3K27me3 in several organs—those containing germline cells (the gonad and oocyte), the head ganglion, and the intestine—of the naïve or *P. aeruginosa* PA14-trained mutants, was decreased compared to that in the organs of the wild-type N2 naïve worms or *P. aeruginosa* PA14-trained control worms. Compared to that in *P. aeruginosa* PA14-trained N2 worms, the acquired robust increase in H3K27me3 signal was almost completely blocked by the mutation of *vit-1* and *nrf-6* (Fig. 3a, b). We also immunostained the *P. aeruginosa* PA14-trained worms with mutations in *vit-1* and *nrf-6* using an antibody against H3K27ac under the same experimental conditions. As expected, H3K27ac levels exhibited the exact opposite pattern (Extended Data Fig. 3a, b) in the organs containing germline cells and other tissues. Nevertheless, the IF staining experiments supported the notion that Vit proteins and their transporter mediate the establishment of acquired H3K27me3 and H3K27ac in the germline of parental worms trained with *P. aeruginosa* PA14.

**Fig. 3.**
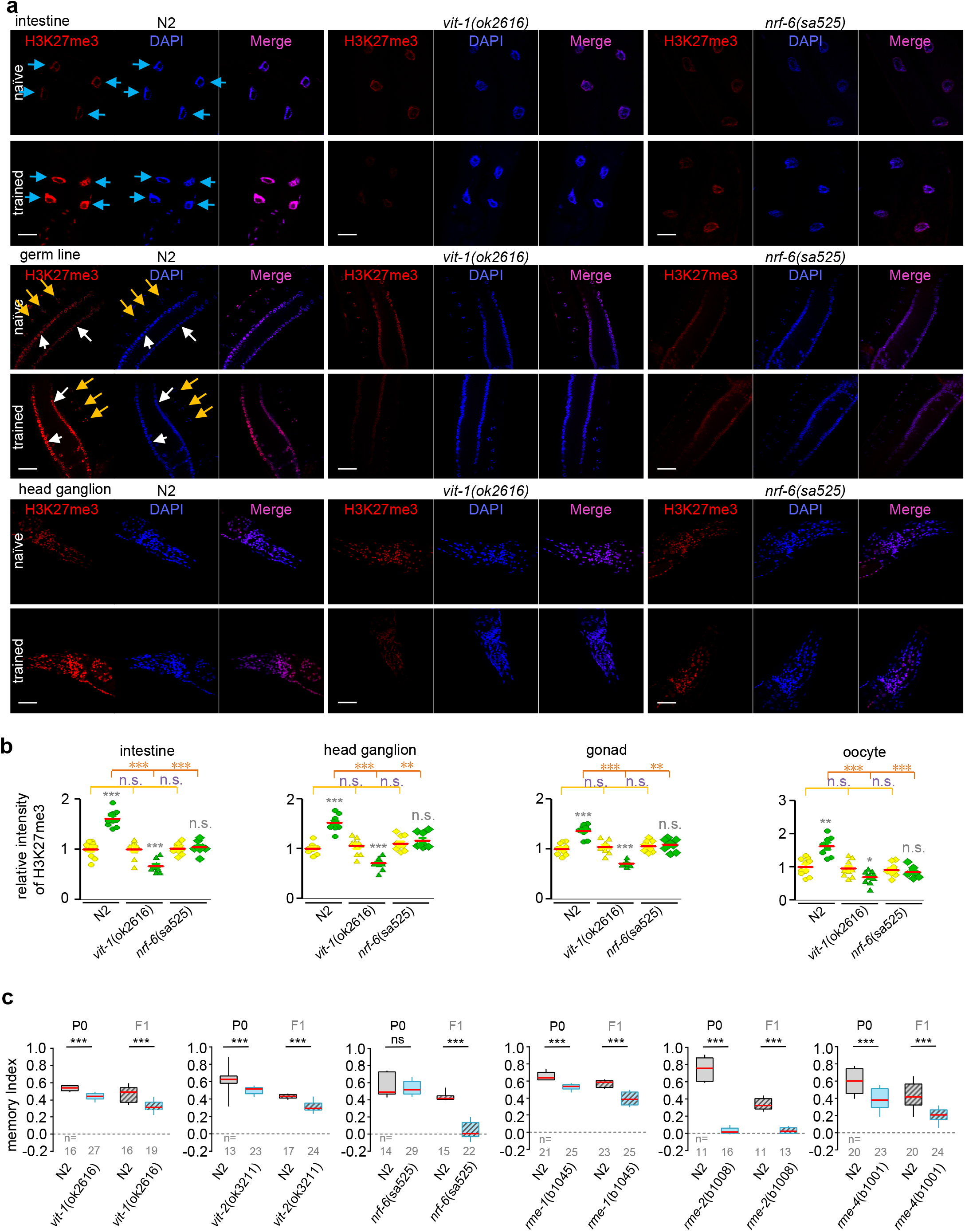
Vitellogenins and their transporters are required to establish H3K27me3 in the parental germline and transgenerational inheritance. **a**, *P. aeruginosa* PA14-induced global increase in H3K27me3 in the intestine, head ganglion and germline in *N2* animals was abolished in both worms expressing the *vit-1 (ok2616)* mutant and those expressing the *nrf-6 (sa525)* mutant. Representative worms were fixed and immunostained with 4’,6-diamidino-2-phenylindole (DAPI) (blue) and antibody against H3K27me3 (anti-H3K27me3) (red). The merged signals are shown in pink. Blue, white and yellow arrows indicate the intestine, gonad and oocyte, respectively. **b**, Quantification of H3K27me3 in the intestine, head ganglion, gonad and oocyte in P0 adult *N2*, *vit-1 (ok2616)* mutant and *nrf-6 (sa525)* mutant animals. n≥8 worms analyzed, means ± SEMs; p-values were generated by one-way ANOVA with the Dunnett correction (n.s., not significant; *p < 0.05, **p < 0.01, and ***p < 0.001). **c**, *P. aeruginosa* PA14-induced transgenerational memory in P0 and F1 worms was blocked in the presence of mutations in genes associated with yolk proteins *(vit-1 (ok2616), vit-2 (ok3211)),* yolk transporter *(nrf-6 (sa525))* and yolk receptors *(rme-1 (b1045), rme-2 (b1008), rme-4 (b1001)).* Filled columns on the right with slanted lines indicate F1 animals, and the two filled columns on the left represent P0 animals. Blue squares represent mutant animals; p-values were generated by one-way ANOVA with the Dunnett correction (n.s., not significant; ***p < 0.001).

Subsequently, we examined whether the loss of yolk proteins or their transporter would also affect the acquired behavior of the *P. aeruginosa* PA14-trained animals. Worms expressing mutants of *vit-1* and *vit-2,* the transporter *nrf-6,* the Vit receptor *rme-2,* and the Vit regulators *rme-1* and *rme-4* were examined and compared with *P. aeruginosa* PA14-trained wild-type P0 control and offspring worms. Compared to control worms, *P. aeruginosa* PA14-trained worms expressing mutants of both *vit-1 (ok3211) and vit-2 (ok3211)* exhibited a more consistent and significant loss of memory behavior in the parental animals and offspring (Fig. 3c). A similar pattern in terms of the loss of transgenerational behavior was observed in worms expressing *rme-1 (b1045), rme-2 (b1008),* and *rme-4 (b1001)* mutants (Fig. 3c). Among the trained mutant worms, the mutation of *nrf-6 (sa525)* seemed to induce a more drastic loss of memory in offspring than the parental (P0) animals, which was possibly due to the increased failure to transmit Vits to oocytes from the intestine under the mutational background (Extended Data Fig. 3c)^17^.

To further understand the mechanism by which Vits/yolk proteins modulate acquired transgenerational epigenetic modifications in the germline, we performed an immunoprecipitation (IP) experiment using transgenic adult worms expressing vit-2-GFP^17^. Control worms and worms expressing vit-2-GFP were therefore collected, lysed, and immunoprecipitated with GFP-conjugated agarose beads (Fig. 4a). The IP products were collected and subjected to mass spectrometry analysis. More than 150 proteins were found to be physically associated with vit-2 (Table S1). Further protein-protein interaction network analysis subdivided the identified proteins into 6 groups based on their function: enzyme modifiers, nicotinic acetylcholine receptors, catalytic proteins, translational and transcriptional regulators, and proteins required for cellular and metabolic processes (Fig. 4b).

**Fig. 4.**
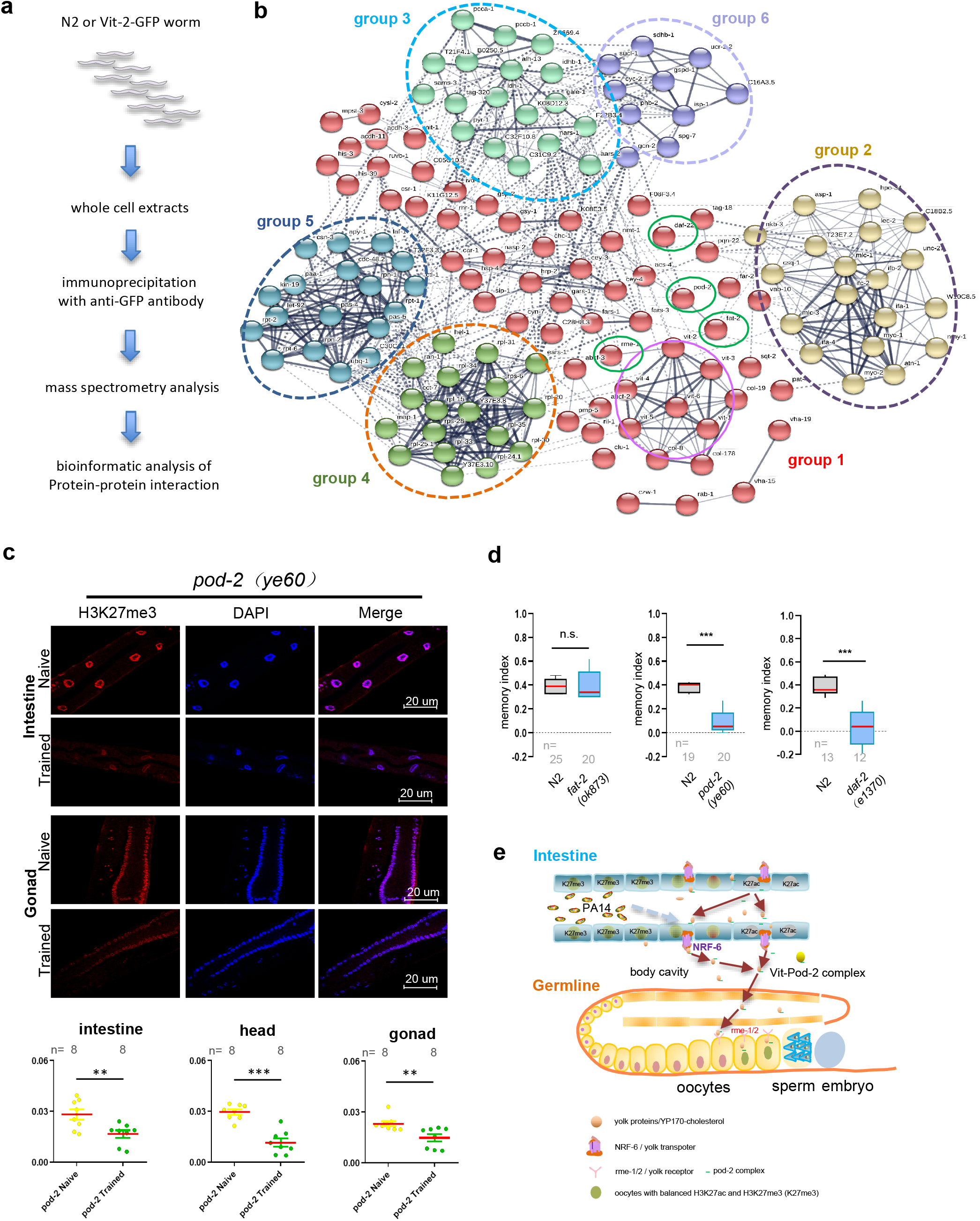
pod-2 interacts with the vitellogenin protein and mediates the acquisition of H3K27me3. **a**, Schematic illustration of the mass spectrometry analysis. **b**, An illustration of interaction networks between the newly identified proteins associated with vit-2. Six groups of proteins are indicated by dotted colored circles. Group 1 includes proteins with catalytic activity and protein binding; group 2, nicotinic acetylcholine receptors; group 3, proteins associated with transcription regulation; group 4, proteins with translation activity; group 5, protein-modifying enzymes; and group 6, proteins associated with cellular and metabolic processes. The selected interacting proteins for further analysis, including pod-2, fat-2 and daf-22, are circled in green. The names of the proteins identified are also indicated next to the colored crystal droplet. **c**, The *P. aeruginosa* PA14-induced increase in H3K27me3 in the intestine and germline of *N2* animals was abolished by the *pod-2 (ye60)* mutation but not the *fat-2 (ok873)* or *daf-22 (ok693)* mutation. Representative worms were fixed and immunostained with 4’,6-diamidino-2-phenylindole (DAPI) (blue) and antibody against H3K27me3 (anti-H3K27me3) (red). The merged signals are shown in pink. Day-1 worms were used for the analysis. Quantification of H3K27me3 in the native and trained intestine, gonad and head ganglion of *pod-2 (ye60)* mutant animals. n≥8 worms analyzed, means ± SEMs; p-values were generated by one-way ANOVA with the Dunnett correction (n.s., not significant; *p < 0.05, **p < 0.01, and ***p < 0.001). **d**, *P. aeruginosa* PA14-induced avoidance memory was blocked by the mutation of pod-2 (*ye60*). Filled columns on the left represent N2 animals. Blue filled columns on the right represent mutant or RNAi animals for pod-2, fat-2 or daf-2; p-values were generated by one-way ANOVA with the Dunnett correction (n.s., not significant; ***p < 0.001). **e**, Schematic model of yolk and pod-2 proteins involved in the establishment of transgenerational epigenetic modifications in the parental female germline. Vits/yolk protein complexes containing pod-2, secreted from the intestine and transported to eggs via the body cavity, maintain the proper level of histone acetylation and methylation in the germline. Under chronic exposure to a pathogenic bacterium, the transportation of vit-pod-2 complexes to eggs is drastically enhanced, resulting in a higher consumption of acetyl CoA and a dramatic reduction in H3K27ac. As a consequence, the level of H3K27me3 is significantly increased in oocytes and somatic cells, serving as a transgenerational imprint in the parental germline.

To investigate whether specific protein partners of vit-2 are involved in establishing the acquired increase in H3K27me3 and decrease in H3K27ac in the germline of *P. aeruginosa* PA14-trained animals, several candidate proteins were selected for further testing. Among the selected proteins, pod-2 is an acetyl-CoA carboxylase, the ratelimiting enzyme in the biosynthesis of fatty acids, and serves as the source of the acetyl group for protein acetylation. The loss of pod-2 or orthologs increased the histone acetylation level of H3K27ac^18,19^. Notably, depletion of pod-2 in worms also downregulated S-adenosylmethionine synthetase, which produces SAM, a methyl group donor required for a variety of phospholipid methyltransferase reactions^20–22^. We therefore reasoned that the Vit-2-interacting partner pod-2 may play a direct role in the acquired decrease in H3K27ac and increase in H3K27me3 in the parental worm germline (Fig. 2). IF experiments were performed to test this hypothesis. Indeed, the increase in H3K27me3 were significantly blocked in the *P. aeruginosa* PA14-trained pod-2 mutant germline when compared to the germline of *P. aeruginosa* PA14-trained control animals (Fig. 4c; Fig. S4). No similar changes in acquired H3K27me3 or H3K27ac modification of fat-2, a stearoyl-CoA desaturase family member, or daf-22, a homologue of the human sterol carrier protein SCPx, were required to catalyze the final step of peroxisomal fatty acid β-oxidation^23^ (Fig. 4C and Extended Data Fig. 4a-c). As expected, parental animals expressing mutant pod-2 exhibited a tremendous loss in memory behavior (Fig. 4d). Consistent with this finding, the insulin/IGF-1-like receptor gene *Daf-2 (e1370)* mutation, known to regulate animal longevity, which causes reduced transcription of the pod-2 partner *Vits^24^,* showed a similar loss of avoidance behavior (Fig. 4).

Environment-induced intergenerational/transgenerational maternal inheritance has been reported in a number of mammalian studies, including the agouti locus^25^, and under a number of conditions, including fetal malnutrition-induced symptoms^26^ and the in utero passage of photoperiod information^27^ in mammals. However, the mechanism driving acquired intergenerational and transgenerational epigenetic changes in the female germline of animals is unclear. Studies in flies and worms have suggested that H3K27me3 in the germline can be transmitted to the next generation^28–30^. This study further shows that the lipoprotein Vits produced in the intestine, together with the acetyl-CoA carboxylase pod-2, serve as a key mediator (Fig. 4e, and Supplementary Table 1) for the establishment of acquired H3K27me3 and H3K27ac in the germline of parental animals. The finding that Vits/yolk proteins and pod-2 regulate the dynamics of H3K27ac and H3K27me3, key epigenetic regulators of various cancers^12, 13, 31^, may also provide an important drug target for certain gut microbiota- or diet-induced intergenerational and transgenerational health problems, such as maternally induced defects of the fetus, diet-induced cancer, diabetes, and mental/behavioral disorders, in humans.

## Methods

### Cultivation of the worms

Worms were grown at 20°C on nematode growth medium (NGM) plates seeded with bacteria *(E. coli* OP50) as a food source. Embryos were collected by hypochlorite bleaching of gravid adult animals and grown at 20°C.

### Binary choice assay

To prepare testing plates, 20 μl of a fresh overnight OP50 and *P. aeruginosa* PA14 bacterial suspension was seeded on a 10-cm NGM round plate that was air-dried for 2 hours at room temperature. Adult hermaphrodites were washed from their growth plate with M9 buffer and rinsed twice, after which approximately 100 animals were placed in the middle of each 10-cm test plate, equidistant from the bacterial lawns^4^. **Training assay.** Naïve worms were cultivated on a standard 6-cm NGM plate seeded with OP50 bacteria. For 72 hours of training, 200 μl of a suspension of fresh *P. aeruginosa* PA14 was spread on the middle of the plate to form a wide barrier lawn, and 50 μl of fresh OP50 suspension was dropped on the plate to form a small lawn on one side of the *P. aeruginosa* PA14 barrier. The training plates were incubated at 26°C for 48 hours before use^1, 2^. Embryos were collected by bleaching and hatching at 20°C on the other side of the *P. aeruginosa* PA14 barrier on the 72-hour training plates. After the embryos had hatched, 72 hours later, day-1 adult worms were collected by washing with M9 buffer three times and then dropped on 10-cm test plates to perform the binary choice assay.

### Transgenerational memory test

F1 generation embryos were collected by hypochlorite bleaching of the P0 generation of the naïve group and 72-hour-trained group (both from test plates on the OP50 sides) immediately after the binary choice assay. F1 generations from the naïve group and trained group were cultivated on standard 6-cm NGM plates seeded with OP50 until day-1 adults were obtained. The F1 generation day-1 adult worms were washed with M9 buffer three times and then placed on 10-cm test plates to perform the binary choice assay. Similar procedures were applied to the F2, F3, F4 and F5 generations of naïve animals.

### Immunostaining

Worms were collected in M9 medium and centrifuged briefly. The worms were then fixed in fixing solution (1x PBS, pH 7.4, 4% paraformaldehyde) at 4°C for 30 min and incubated in 5% 2-mercaptoethanol at 37°C overnight after washing three times in washing solution (1× PBS, pH 7.4, 0.5% Triton X-100). The worms were washed in washing solution 3 times and incubated in collagenase at 37°C for 6 hours. The worms were then washed in washing solution and blocked in PBST (1× PBS, pH 7.4, 1% BSA, 0.5% Triton X-100) for 2 hours. After the blocking solution was removed, the worms were incubated overnight at 20°C with primary antibodies diluted in PBST. The anti-H3K27me3 antibody (Abcam, ab6002) was used at a 1:100 dilution, and the anti-H3K27me3 antibody (Millipore, 07-449), anti-H3K27ac antibody (Abcam, ab4729) and anti-H4K16ac antibody (Abcam, ab109463) were used at a 1:200 dilution. The worms were washed 3 times for 10 min each in washing solution and then incubated with secondary antibody (1:800 in 1% BSA) for 4 hours at 20°C. Alexa Fluor 568-conjugated goat anti-mouse (Life Technology A-11004) was used to detect the H3K27me3 antibody. The worms were then washed 4 times in washing solution for 7 min each and mounted with ProLong Gold antifade reagent with DAPI.

### Imaging

Images of fixed worms were acquired using an alpha Plan-Apochromat 100x/1.46 oil immersion objective on a Zeiss confocal LSM 880 Axio Observer microscope. Images were processed by ZEN 2.1 SP3 software. Live worms were anesthetized using 10 mM NaN3 on 2% agarose pads as previously described^32^, and Z-stack imaging was performed on a Zeiss Axio Imager. An M2 microscope with Apotome under a 20x objective was used. Images were acquired with a Hamamatsu CCD camera (ORCA-Flash4.0 V2), and maximum-intensity projections were built with Zeiss ZEN 2.3 pro. Yolk imaging was performed on a Zeiss Observer 7 microscope under a 100x objective. Images were acquired with a photometrics CCD (PRIME) camera and processed by Zen2.3 pro software. Single-cell images were acquired with a Zeiss V16 microscope using ImageJ 1.51d.

### IP and mass spectrometry

Transgenic worms expressing vit-2-GFP and control N2 worms were collected with 15 ml of M9 buffer (0.6% Na2HPO4, 0.3% KH_2_PO_4_, 0.5% NaCl and 1 M MgSO_4_·7H_2_O) by centrifugation at 2,000 rpm for 1 min. The supernatant was removed, and 1 ml of ice cold lysis buffer (25 mM Tris-HCl (pH 7.5), 100 mM NaCl, 1 mM EDTA, 0.5% NP-40, 1 mM PMSF, 1 mM Na_3_VO_4_, 1 μg/ml Pepstatin-A, 10 mM NaF and protease inhibitor cocktail) was added to the worms, which were incubated on ice for 30 min. The worms were frozen in liquid nitrogen and then thawed at room temperature, after which they were further sonicated for 7 sec using 12 W of power 3 times. Samples were rotated at 4 °C by endover-end agitation for 30 min and centrifuged at 13,000 rpm for 30 min at 4 °C. The supernatant was then transferred into a 1.5-ml tube. The protein concentration was measured using a BCA protein assay kit (#53225, Thermo Fisher). Four milligrams of total protein was mixed with 20 μl of GFP-conjugated agarose beads (#gta-20, Chromotek) equilibrated by washing in lysis buffer. The mixed samples were incubated at 4 °C with agitation overnight. After incubation, the beads were washed 3 times using 1 ml of ice cold lysis buffer each time, and the samples were incubated with agitation at 4 °C for 5 min between washes. Proteins were eluted with 40 μl of SDS sample buffer and boiled for 8 min. The samples were then subjected to SDS-PAGE and stained with Coomassie Brilliant Blue. The mixed protein bands were collected and subjected to mass spectrometry (Q Exactive^TM^ Plus, Thermo) analysis by Jingjie PTM Bio (Hangzhou, China).

### RNA extraction

After the behavioral test, N2 worms were collected from the naïve and trained (the OP50 side and *P. aeruginosa* PA14 side of the test plate, respectively) groups in M9 buffer. RNA was extracted from the worms using the standard TRIzol (Takara) method, followed by chloroform precipitation.

### GO analysis

The R package ClusterProfiler^33^ was used to implement the GO overrepresentation test^34^ and gene set enrichment analysis (GSEA)^35^ with the Bioconductor annotation package^36^ of *C. elegans.* Visualization of the results of GO term analysis was conducted with ggplot2^37^ v3.2.0 (for dot plots and Cleveland dot plots) and GOplot ^38^ v1.0.2 (for “golden eye” plots).

### Data availability and associated accession codes

The accession number for the sequencing data reported in this paper is GEO: GSE 138059.

## Supporting information

Supplementary Resource Table

## Acknowledgments

We thank all the members of the W.L. and F.L.S. laboratories for their discussion and constructive suggestions regarding the project.

## Funding

This work was supported by the “National Key Research and Development Program of China” (grant No.: 2017YFA0103301, 2016YFA0100403), a research grant from the Tsingtao Institute for Advanced Research, Tongji University, and a grant supported by the National Natural Science Foundation of China (grant No: 31330043), with partial financial support from the Alpha Institute of Natural Medicine, Tongji University.

## Author contributions

F.L.S. and W.L. designed the experiments and contributed to data analysis, discussion, and writing. D.P., C.W., K.L.L., Z.X.G., H.W.W., Q.Y.L., H.Y.S., Y.Y.J., Q.F., Q.Z., L.Z., L.J., H.H.C., H.M.W., X.M.Y., J.F.C., F.S. and H.Y.S. contributed to the experimental work and discussion. Y.H.L. H.Y.T., J.S., C.Z.J., and X.W.L. contributed to the bioinformatic analysis.

## Competing interests

The authors declare no competing interests.

## Supplementary materials

Supplementary Materials and Methods, resource data, figure legends, references and associated codes will be available in the online version of the paper.

## Data and materials availability

All data needed to understand and acquire the conclusions of this study are included in the text, figures, and supplementary materials.

---

**Extended Data Fig. 1.**
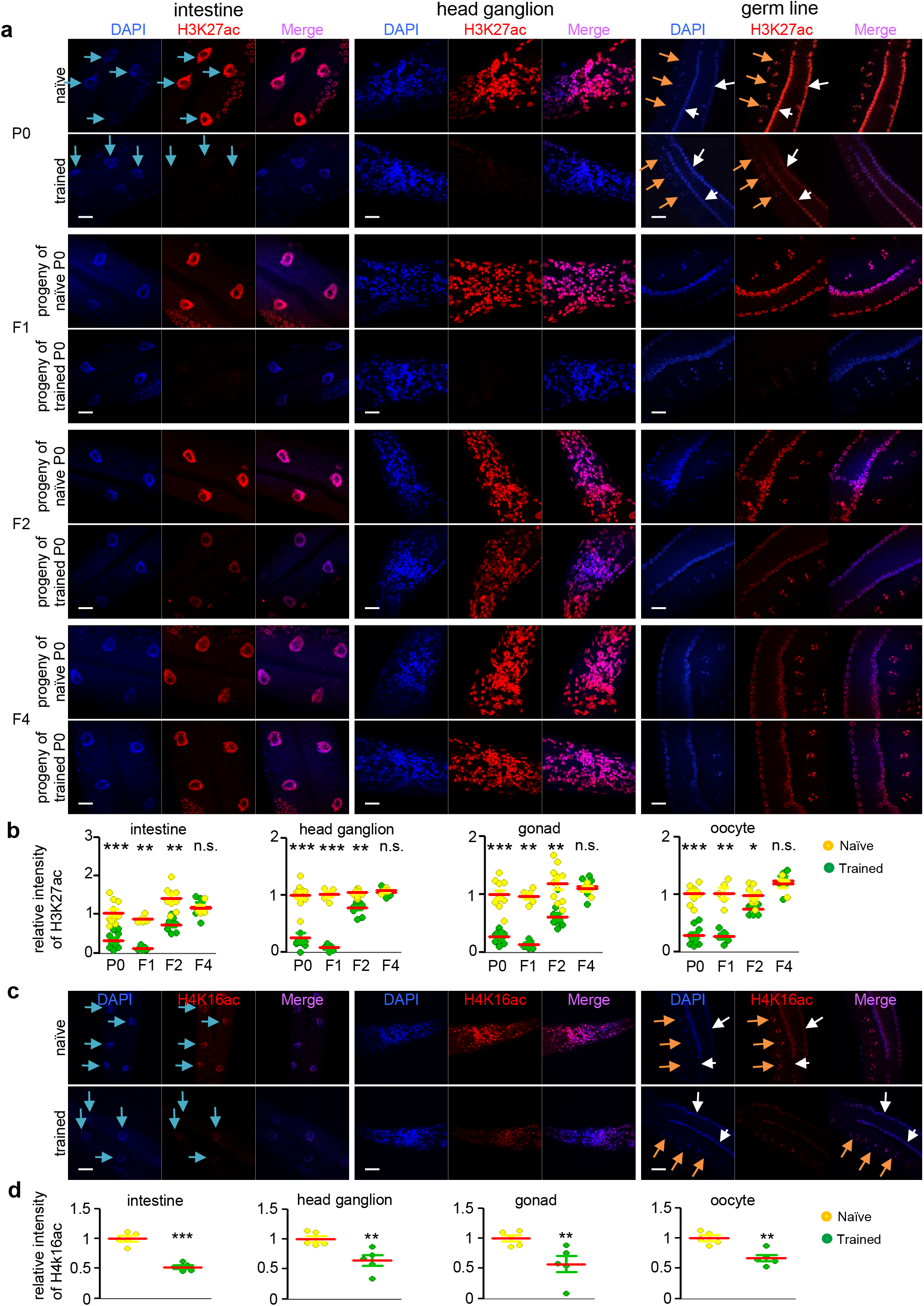
Screening of hallmarks for pathogenic bacterium-induced parental germline and transgenerational inheritance. **a**, Illustration of *P. aeruginosa* PA14-induced repression of the global levels of H3K27ac in the intestine, head ganglion and germline in P0 adult animals and their F1 and F2 descendants but not in their F4 descendants. Representative worms were fixed and immunostained with 4’,6-diamidino-2-phenylindole (DAPI) (blue) and antibody against H3K27ac (anti-H3K27ac) (red). The merged signal is shown in pink. **b**, Quantification of the mean level of nuclear H3K27ac in the intestine, head ganglion, gonad and oocyte in P0 adult animals and their F1, F2 and F4 descendants. Means ± SEMs; p-values were generated by one-way ANOVA with the Dunnett correction (ns, not significant; *p < 0.05, **p < 0.01, and ***p < 0.001). Blue, white and yellow arrows indicate the intestine, gonad and oocyte, respectively. **c**, *P. aeruginosa* PA14 induced a reduction in global levels of H4K16 histone acetylation (H4K16ac) in the intestine, germline, and head and tail ganglia in P0 adult animals. Representative worms were fixed and immunostained with DAPI (blue) and antibody against H4K16ac (anti-H4K16ac) (red). **d**, Quantification of the mean nuclear H4K16ac signal intensity in the intestine, head ganglion, gonad and oocyte in P0 adult animals. Means ± SEMs; n≥5; p-values were generated by one-way ANOVA with the Dunnett correction (ns, not significant; *p < 0.05, **p < 0.01, and ***p < 0.001).

**Extended Data Fig. 3.**
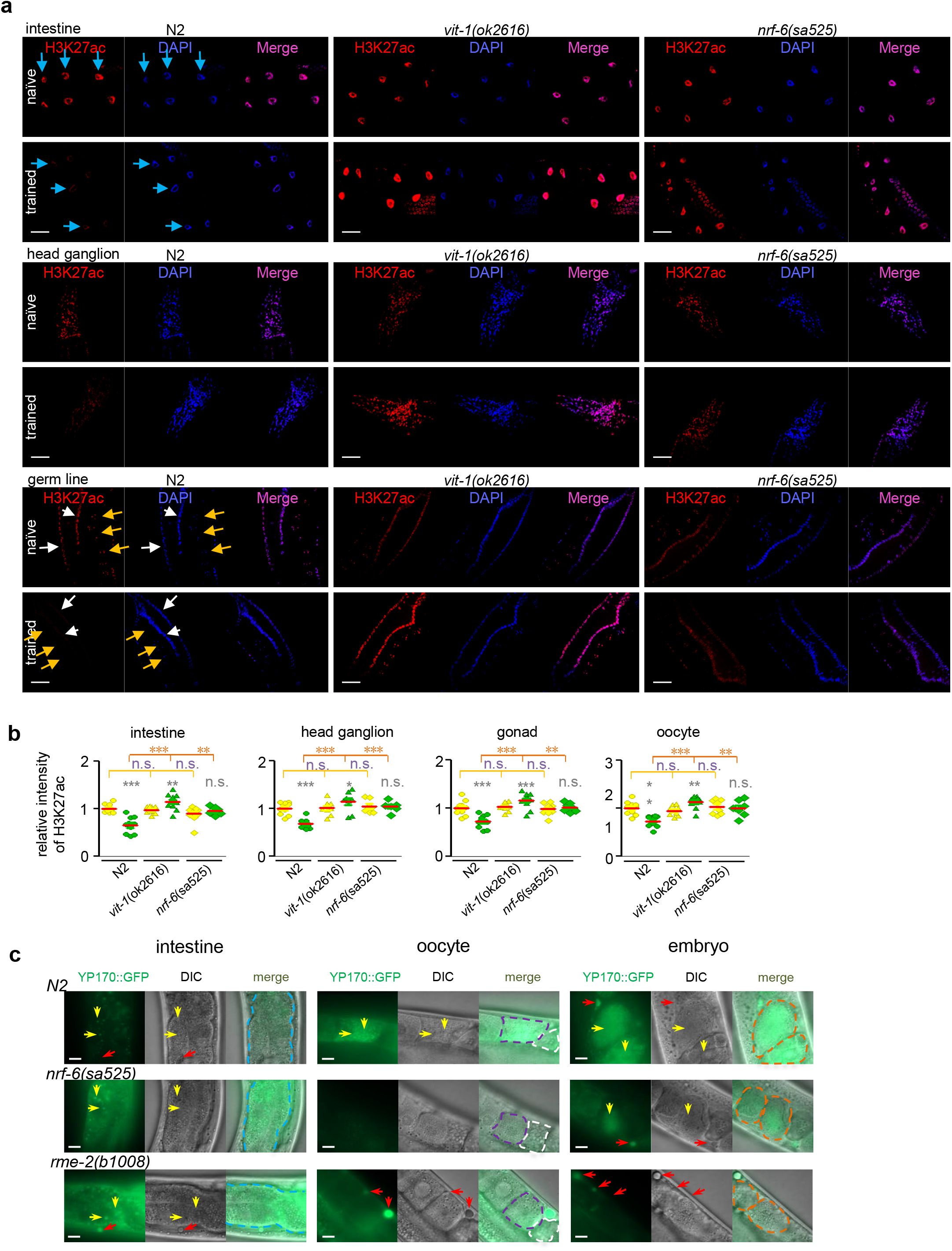
Vitellogenins and their transporters are required for H3K27ac in the parental germline. **a**, The *P. aeruginosa* PA14-induced global reduction in H3K27 levels in the intestine, head ganglion and germline of N2 animals was abolished in *vit-1 (ok2616)* mutant but not in *nrf-6 (sa525)* mutant worms. Representative worms were fixed and immunostained with 4’,6-diamidino-2-phenylindole (DAPI) (blue) and antibody against H3K27ac (anti-H3K27ac) (red). The merged signals are shown in pink. White and yellow arrows indicate gonads and oocytes, respectively. **b**, Quantification of H3K27ac in the intestine, head ganglion, gonad and oocyte in P0 adult N2, *vit-1 (ok2616)* mutant and *nrf-6 (sa525)* mutant animals. N≥8 worms analyzed, means ± SEMs; p-values were generated by one-way ANOVA with the Dunnett correction (ns, not significant; *p < 0.05, **p < 0.01, and ***p < 0.001). **c**, Illustration of yolk protein expression, as indicated by the YP170 GFP-tag levels in different organs, namely, the intestine, oocyte and embryo, of wild-type N2 worms and N2 worms with a background of *nrf-6 (sa525)* and *rme-2 (b1008)* mutation. The yellow arrows indicate yolk proteins, and the red arrows indicate free yolk in the body cavity. The blue dotted circles indicate the intestinal cells, the white dotted circles indicate the spermatheca, the purple dotted circles indicate the oocyte, and the orange dotted circles indicate the two embryos.

**Extended Data Fig. 4.**
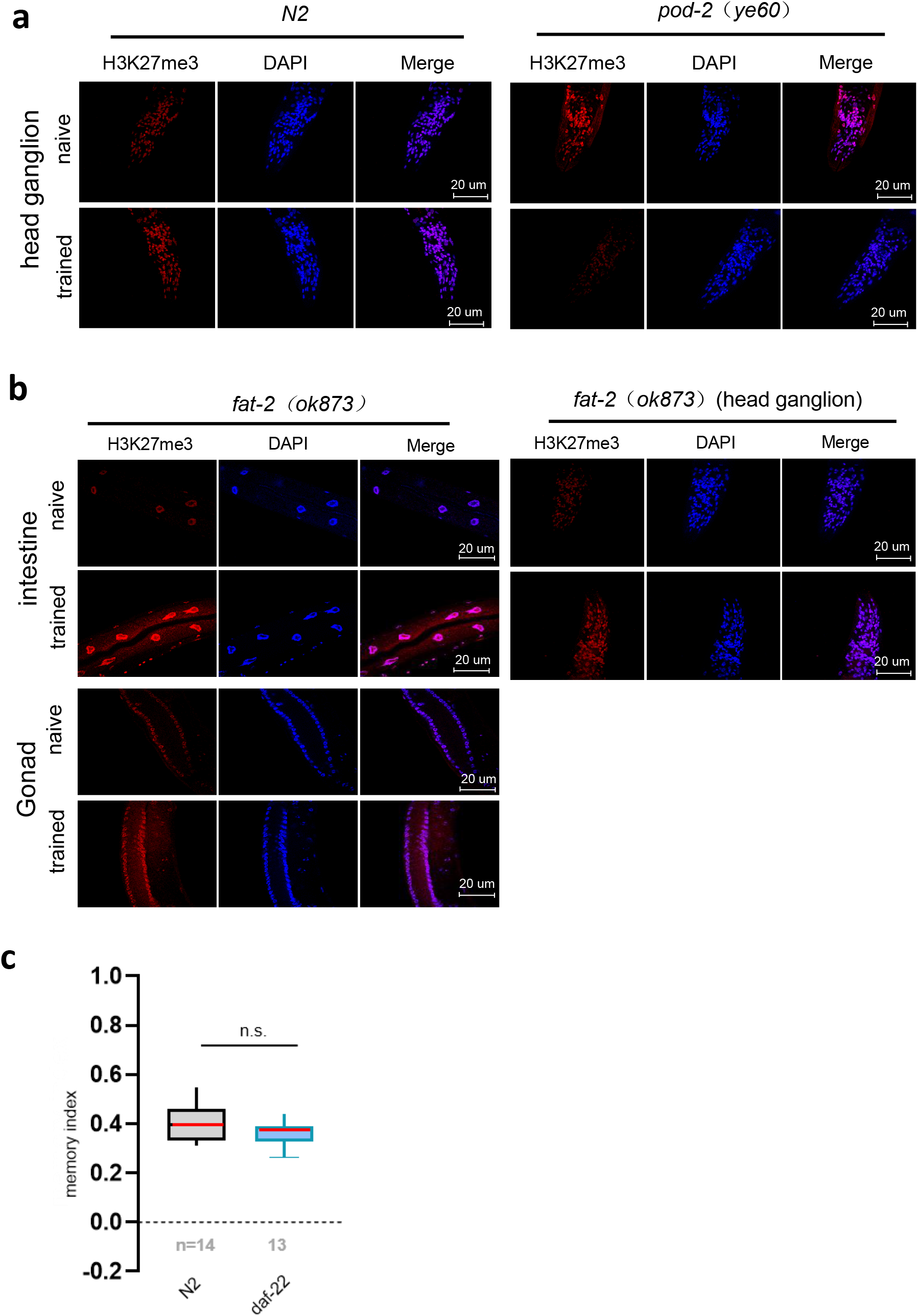
pod-2 mediates the acquisition of H3K27me3. **a**, The *P. aeruginosa* PA14-induced increase in H3K27me3 levels in the head ganglion of N2 animals was abolished in the *pod-2 (ye60)* mutant worms but not in the *fat-2 (ok873)* mutant worms. **b**, Representative worms were fixed and immunostained with 4’,6-diamidino-2-phenylindole (DAPI) (blue) and antibody against H3K27me3 (anti-H3K27me3) (red). The merged signals are shown in pink. Day-1 worms were used for the analysis. **c**, *P. aeruginosa* PA14-induced avoidance memory by the depletion of *Daf-22.* Filled columns on the left represent N2 animals. Blue filled columns on the right represent animals in which *daf-22* was knocked down via RNAi; p values were generated by one-way ANOVA with the Dunnett correction (n.s., not significant).

## Notes

### Competing Interest Statement

The authors have declared no competing interest.

