## Supplementary Resource Table for "The Establishment of Transgenerational Epigenetic Inheritance in the *C. elegans* Germline is Mediated by Lipid Metabolism"

| Reagent or Resource | Source | Identifier |
| --- | --- | --- |
| Antibodies | | |
| rabbit anti-H3K27me3 antibody | Millipore | 07-449 |
| mouse anti- H3K27me3 antibody | abcam | ab6002 |
| rabbit anti- H4K16ac antibody | abcam | ab109463 |
| rabbit anti-H3K27ac antibody | abcam | ab4729 |
| Alexa Fluoro 568 goat anti-mouse | Life Technology | A-11004 |
| Cy3-anti-rabbit | Jackson Immunoresearch | 711-165-152 |
| Bacterial and Virus Strains | | |
| *E. coli* OP50 | Caenorhabditis Genetics Center | CGC *E. coli* Strain: OP50 |
| *P. aeruginosa* PA14 | Ausubel Lab, Harvard Medical School | N/A |
| Chemicals, Peptides, and Recombinant Proteins | | |
| YEAST EXTRACT | OXOID | LP0021; lot#1213550-02 |
| TRYPTONE | OXOID | LP0042; lot#1939097 |
| PEPTONE No.2 | OXOID | LP0137; lot#1901147 |
| NaOH | Sigma-Aldrich | S5881; CAS: 1310-73-2; lot#BCBH5016V |
| MgSO_4_ | Sigma-Aldrich | M7506; CAS: 7487-88-9; lot#WXBC7784V |
| Na_2_HPO_4_·2H_2_O | Sigma-Aldrich | 71645; CAS:10024-24-7; lot#BCBR5790V |
| KH_2_PO_4_ | VETEC | V900041; CAS:7778-77-0; lot#WXBC4661V |
| CaCl_2_ | VETEC | V900266; CAS: 10043-52-4; lot#WXBB1152VV |
| NaCl | VETEC | CAS: 7647-14-5; lot#MKBR1837V |
| RNAiso Plus | Takara | CAT#9109 |
| paraformaldehyde | Sigma-Aldrich | CAS: 30525-89-4  lot # WXBC2903V |
| 2-mercaptoethanol | Amresco | CAS: 60-24-2  lot # 2674C228 |
| Bovine Serum Albumin | VETEC | CAS: 9048-46-8  Lot#WXBC2903V |
| KCl | Sigma-Aldrich | 746436; CAS:7447-40-7; lot#SLBR9127V |
| HEPES | Sigma-Aldrich | H4304; CAS:7365-45-9; lot#100M5418V |
| MgCl_2_·6H2O | Sigma-Aldrich | M2393; CAS:7791-186; lot#081M00011V |
| CaCl_2_·2H2O | Sigma-Aldrich | C3881; CAS:10035-04-8; lot#SLBB9179V |
| 1,4-Dithio-DL-threitol | Sinopharm Chemical Reagent Co.,Ltd | CAS:3483-12-3; lot#63002631 |
| SDS | Biodee | DE-L5750S |
| Collagenase | Sigma-Aldrich | CAS: C0130-500MG |
| Potassium chloride | Sigma-Aldrich | 746436; CAS: 7447-40-7 |
| Qubit™ dsDNA HS Assay Kit | Invitrogen | Q32854 |
| KAPA HiFi HotStart ReadyMix (500 rxn) | KAPA | kk2602 |
| KAPA Hyper Prep Kits with PCR Library Amplification/Illumina series (96 reactions) | KAPA | kk8504 |
| DNA Clean & Concentrator™-5 (ZYMO kit) | ZYMO | D4014 |
| NEBNext® Multiplex Oligos for Illumina® (Index Primers Set 1) | NEB | E7335L |
| Magnesium chloride hexahydrate | VETEC | V900020; CAS: 7791-18-6 |
| Calcium chloride dihydrate | Sigma-Aldrich | C3881; CAS: 10035-04-8 |
| Sodium chloride | VETEC | V900058; CAS: 7647-14-5 |
| HEPES | Sigma-Aldrich | H4034; CAS: 7365-45-9 |
| Triton™ X-100 | Sigma-Aldrich | T9284; CAS: 9002-93-1 |
| Betaine | SAFC | 61962; CAS: 107-43-7 |
| RNaseZap™ RNase Decontamination Solution | Invitrogen | AM9780 |
| UltraPure™ DNase/RNase-Free Distilled Water | Invitrogen | 10977023 |
| Ambion™ RNase Inhibitor, cloned, 40 U/µ | Invitrogen | AM2684 |
| dNTP Mix (10 mM each) | Thermo Scientific | R0193 |
| SuperScript™ II Reverse Transcriptase | Invitrogen | 18064071 |
| Dynabeads™ MyOne™ Streptavidin C1 | Invitrogen | 65001 |
| AMPure XP beads(60ml) | Beckman | A63881 |
| DAPI | Sigma-Aldrich | D9542 |
| Tween 20 | General-Reagent | G89190B; CAS: 9005-64-5 |
| Trichostatin A (TSA) | selleck | CAT# S1045 |
| GSK343 | selleck | CAT#S7164 |
| Deposited Data | | |
| Raw and analyzed data | This paper | GEO: GSE 138059 |
| Experimental Models: Organisms/Strains | | |
| *C. elegans*: Strain N2 (wild type) | Caenorhabditis Genetics Center | WB Strain: N2 |
| *C. elegans*: Strain JT525: *nrf-6(sa525)II* | Caenorhabditis Genetics Center | WB Strain: JT525 |
| *C. elegans*: Strain RB1618: *hda-3(ok1991)I* | Caenorhabditis Genetics Center | WB Strain: RB1618 |
| *C. elegans*: Strain RB758: *hda-4(ok518)X* | Caenorhabditis Genetics Center | WB Strain: RB758 |
| *C. elegans*: Strain RB2357: *F41H10.6(ok3203)IV* | Caenorhabditis Genetics Center | WB Strain: RB2357 |
| *C. elegans*: Strain DH1201: *rme-1(b1045)V* | Caenorhabditis Genetics Center | WB Strain: DH1201 |
| *C. elegans*: Strain DH1390: *rme-2(b1008)IV* | Caenorhabditis Genetics Center | WB Strain: DH1390 |
| *C. elegans*: Strain RT362: *rme-4(b1001)X; Pwls23* | Caenorhabditis Genetics Center | WB Strain: RT362 |
| *C. elegans*: Strain RB1982: *vit-1(ok2616)X* | Caenorhabditis Genetics Center | WB Strain: RB1982 |
| *C. elegans*: Strain RB2365: *vit-2(ok3211)X* | Caenorhabditis Genetics Center | WB Strain: RB2365 |
| *C. elegans*: Strain DH1370: *rme-6(b1014)I* | Caenorhabditis Genetics Center | WB Strain: DH1370 |
| *C. elegans*: Strain DH1206: *rme-8(b1023)I* | Caenorhabditis Genetics Center | WB Strain: DH1206 |
| *C. elegans*: Strain BCN9071: *vit-2(crg9070[vit-2::gfp])X* | Caenorhabditis Genetics Center | WB Strain: BCN9071 |
| *C. elegans*: Strain *mes-2 (tm5007)II* | National BioResource Project | NBRP Strain: tm5007 |
| *C. elegans*: Strain wyIs592: *wyIs592[Pser2prom3::GFP;Odr-1::RFP]III* | Zou Lab, Shanghai Tech University | Zou Lab Strain: wyIs592 |
| *C. elegans*: Strain YV444: *ivyEx148[vit-2(crg9070[vit-2::gfp])X,Pser-2prom3::mcherry]* | This paper | N/A |
| *C. elegans*: Strain YV446: *nrf-6(sa525)II; vit-2(crg9070[vit-2::gfp]X* | This paper | N/A |
| *C. elegans*: Strain YV447: *rme-2(b1008)IV; vit-2(crg9070[vit-2::gfp]X* | This paper | N/A |
| Recombinant DNA | | |
| Ppd95.75-Pser-2prom3::mcherry plasmid | This paper | N/A |
| Software and Algorithms | | |
| FastQC v0.11.8 | Andrews S. , 2010. | http://www.bioinformatics.babraham.ac.uk/projects/fastqc |
| MultiQC v1.0 | *Ewels P.,et al. 2016* | https://multiqc.info/ |
| Kallisto v0.46.0 | Bray N. et al., 2016 | https://pachterlab.github.io/kallisto/ |
| UMI-tools v0.5.5/v1.0.0 | Smith T. et al., 2017 | https://github.com/CGATOxford/UMI-tools |
| Subread v1.6.4 function featureCounts | Liao Y. et al., 2014 | http://subread.sourceforge.net/ |
| Slueth v0.30.0 | Pimentel H. et al.,2017 | https://pachterlab.github.io/sleuth/ |
| Seurat v3.0.1/v3.0.2 | Stuart et al., 2019 | https://github.com/satijalab/seurat |
| SCDE v1.99.4 | Kharchenko et al.,2019 | http://hms-dbmi.github.io/scde/ |
| CluserProfiler v3.12.0 | Yu G. et al., 2012 | https://bioconductor.org/packages/release/bioc/html/clusterProfiler.html |
| ggplot2 v3.2.0 | Wickham H. et al., 2016 | https://ggplot2.tidyverse.org/ |
| **GOplot v**1.0.2 | Walter et al. 2015 | https://wencke.github.io/ |
| Circos Table Viewer v0.63-9 | Krzywinski, M. et al. 2009 | http://mkweb.bcgsc.ca/tableviewer/ |
| STAR v2.7.1a. | Dobin A. et al., 2013 | https://github.com/alexdobin/STAR |
| Samtools v1.9 | Li H., et al., 2009 | http://samtools.sourceforge.net/ |
| R | R Core Team, 2018 | https://www.r-project.org/ |
| RStudio | RStudio Team, 2015 | https://www.rstudio.com/ |
| Circlize v0.4.6 | Gu Z. et al. 2014 | https://github.com/jokergoo/circlize |
| GraphPad Prism | GraphPad Software | https://www.graphpad.com/scientific-software/prism/ |
| ZEN 2.3 blue edition | Carl Zeiss AG | https://www.zeiss.com/microscopy/int/products/microscope-software/zen.html |
| ImageJ 1.51d | Schneider et al., 2012 | https://imagej.net/Welcome |
| Other | | |
| Wormbase | Kevin L. et al., 2016 | https://wormbase.org/ |
| Wormbase ParaSite BioMart | Howe K. et al., | https://parasite.wormbase.org/biomart/martview/ |
| Database of Ligand-Receptor Partners (DLRP) | Salwinski, L. et al., 2004 | http://dip.doe-mbi.ucla.edu/dip/DLRP.cgi |
| IUPHAR | Harmar,A.J. et al., 2009 | http://www.guidetopharmacology.org/ |
